# Heterogeneous Absorption of Antimicrobial Peptide LL37 in *Escherichia coli* Cells Enhances Population Survivability

**DOI:** 10.1101/313536

**Authors:** Mehdi Snoussi, John Paul Talledo, Nathan-Alexander Del Rosario, Bae-Yeun Ha, Andrej Košmrlj, Sattar Taheri-Araghi

## Abstract

Antimicrobial peptides (AMPs) are broad spectrum antibiotics that selectively target bacteria. Here we investigate the activity of human AMP LL37 against *Escherichia coli* by integrating quantitative, population and single-cell level experiments with theoretical modeling. Our data indicate an unexpected, rapid absorption and retention of a large number of LL37 by *E. coli* cells upon the inhibition of their growth, which increases the chance of survival for the rest of population. Cultures with high-enough cell density exhibit two distinct subpopulations: a non-growing population that absorb peptides and a growing population that survive owing to the sequestration of the AMPs by others. A mathematical model based on this binary picture reproduces the rather surprising behaviors of *E. coli* cultures in the presence of LL37, including the increase of the minimum inhibitory concentration with cell density (even in dilute cultures) and the extensive lag in growth introduced by sub-lethal dosages of LL37.

## Introduction

Antimicrobial peptides (AMPs) are natural amino-acid based antibiotics that are part of the first line of defense against invading microbes in multicellular systems Zasloff, M (2002); Brogden (2005). In humans, AMPs are found in many organs that are in contact with the outside world, including airways, skin, and the urinary tract Hancock and Lehrer (1998); Zasloff, M (2002); Brogden (2005); Jenssen et al. (2006); Ganz (2003); Epand and Vogel (1999). The short sequence of the AMPs (typically <50 amino acids) along with the flexibility in the design and synthesis of new peptides has spurred attention towards understanding the detailed mechanism of AMPs action which can lead to the rational design of novel antibiotic agents Zasloff, M (2002); Brogden (2005); Hancock and Sahl (2006).

A hallmark of the AMPs antibacterial mechanism is the role of physical interactions. AMP’s structures exhibit two common motifs: cationic charge and amphiphilic form Zasloff, M (2002); Brogden (2005). The cationic charge enables them to attack bacteria, enclosed in negatively charged membranes, rather than mammalian cells, which possess electrically neutral membranes. The amphiphilic structure allows AMPs to penetrate into the lipid membrane structures Matsuzaki et al. (1995); Shai (1999); Ludtke et al. (1996); Heller et al. (2000); Taheri-Araghi and Ha (2007); Huang (2000); Yang et al. (2001).

Despite our detailed knowledge on AMP’s interactions with membranes, we lack a comprehensive picture of the dynamics of AMPs in a population of cells. We are yet to determine the extent to which the AMP’s physical interactions disrupt biological processes in bacteria and the degree to which electrostatic forces govern the diffusion and partitioning of AMPs among various cells. Specifically, it was suggested by Matsuzaki and Castanho *et al.* that the density of cells in a culture can alter the activity of AMPs through distributions among different cells Matsuzaki (1999); Melo et al. (2009). We have recently examined the role of adsorption on various cell membranes theoretically Bagheri et al. (2015). Experimental investigations using bacteria and red blood cells by Stella and Wimley groups Savini et al. (2017); Starr et al. (2016) directly demonstrated the decisive role of cell density on the effectivity of antimicrobial peptides.

In this work, we utilize complementary experimental and modeling approaches to understand the population dynamics of AMP’s activity from a single-cell perspective. Like all antibiotic agents, AMPs need a minimum concentration (MIC) to inhibit growth of a bacterial culture. For some antibiotics, including AMPs, the MIC is dependent on the cell density. Often referred to as the “inoculum effect”, this phenomena is a trivial consequence of overpopulated cultures. However, in dilute cultures, MICs have been reported to reach a plateau independent of cell density Savini et al. (2017); Starr et al. (2016); Udekwu et al. (2009); Artemova et al. (2015), unless the cell population becomes so small that stochastic single-cell effects become important Coates et al. (2018).

For a precise measurement of the inoculum effect, we extended microplate assays by ***Wiegand et al.*** (***2008***) to obtain a functional form of the MIC in terms of the initial cell density (the “inoculum size”). Contrary to our expectations, we observed that the MIC for the LL37 peptide (AnaSpec, California) remains dependent on *Escherichia coli* density, even in dilute cultures where the average cell-to-cell distance is above 50 μm, much greater than the average cell dimensions (~1×1×5 μm) Taheri-Araghi et al. (2015). With no direct interactions among the cells and nutrients in excess for all, this dependence suggests that the *effective* peptide concentration is somehow compromised in a cell density dependent manner.

By tracking a dye-tagged version of LL37 peptide, we found that the inhibition of growth of *E. coli* cells was followed by the translocation of a large number of AMPs into the cells cytoplasm, thus reducing the peptide concentration in the culture, which works in favor of other cells. In the sense of such dynamics, MIC refers to a sufficient concentration of AMPs for absorption into all the cells. Below the MIC, peptide is absorbed by only a fraction of cells, leaving an inadequate amount of AMPs to inhibit the growth of remaining cells. We have directly observed that cultures with sub-MIC concentrations of dye-tagged LL37 exhibit a heterogenous population combining non-growing cells containing many LL37 and growing cells without LL37.

## Results

### The MIC increases as a function of cell density

The MIC of an antibiotic may depend on the cell density for various reasons: the distribution of antibiotic molecules among bacteria Udekwu et al. (2009); Clark et al. (2009); Melo et al. (2009); Jepson et al. (2016); or as a result of cellular enzyme secretion (e.g., *β*-lactamase in case of lactame resistant antibiotics Clark et al. (2009); Artemova et al. (2015)); or due to the chemical composition of the culture and regulation of gene expression (for instance in late exponential or stationary phases) Karslake et al. (2016); Artemova et al. (2015).

In this work we focus on dilute cultures where the dependence of MIC on inoculum size reflects the number of antimicrobial molecules either “consumed” or “destroyed” by each individual cell. The MIC for AMPs is in the micromolar range, ~10^14^ AMPs/ml. Early exponential cultures contain ~10^6^ cells/ml, which amounts to the ratio of ~10^8^ AMPs/cell. At such a high ratio, only binding or degradation of AMPs of the same order of magnitude per individual cells can lead to the inoculum effect Starr et al. (2016); Savini et al. (2018).

To map out the functional form of the inoculum effect, we implemented a two-dimensional dilution scheme on a 96-well plate (Fig. 1A). The scheme incorporates a linear dilution of LL37 in columns 7 and 12 followed by a 2/3 dilution series of cells and LL37 over two distinct regions, columns 12 to 8 and 7 to 1. (See SI Figs. S1 and S2 for details of the cell counting and plate preparation). An early exponential *E. coli* culture in rich de1ned media (RDM, Teknova) was diluted to specific cell densities to cover a relatively even distribution of inoculum sizes.

**Figure 1.**
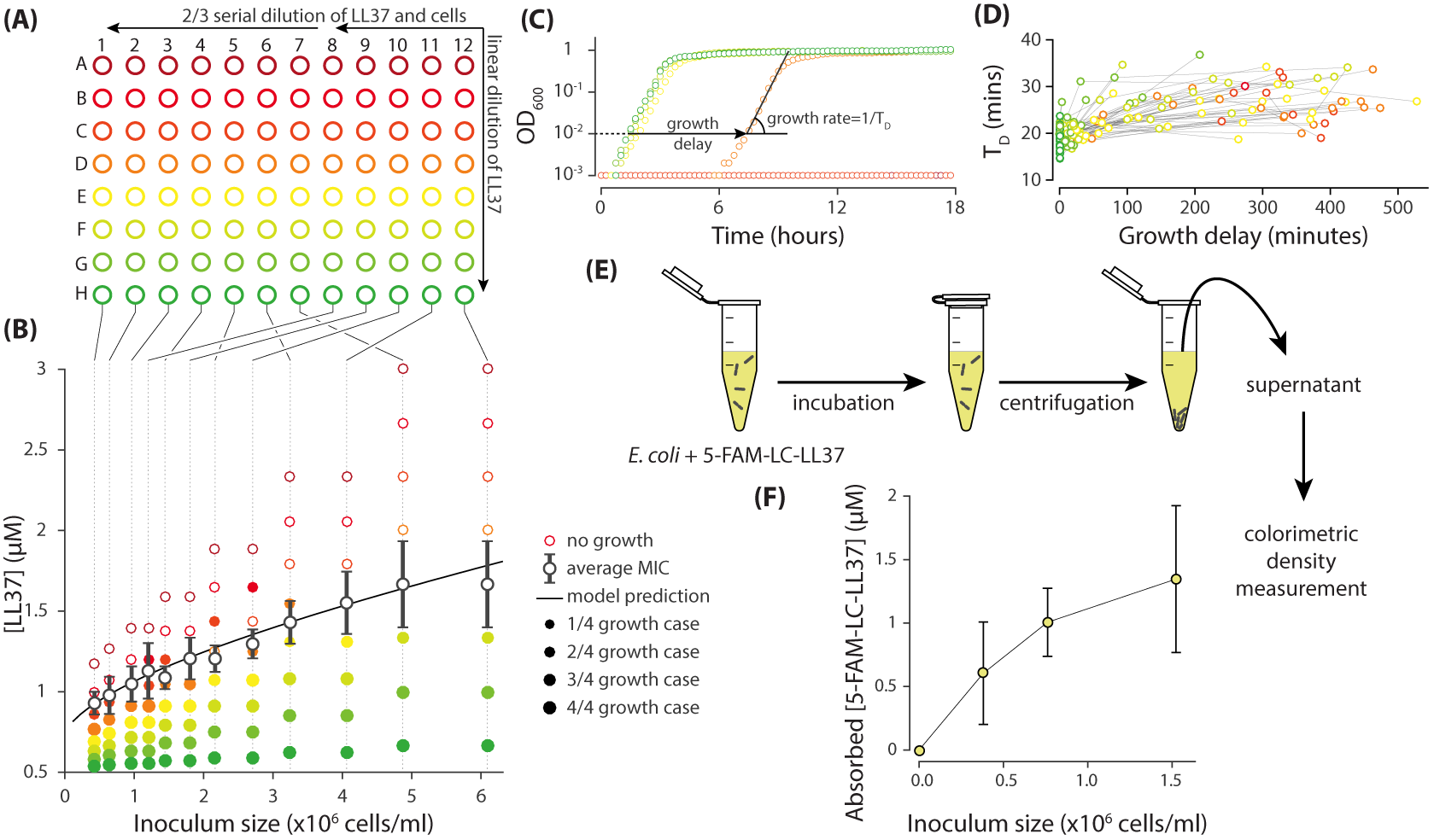
Measurement of the inoculum effect and peptide absorption by *E. coli* cells. (A) A two-dimensional dilution scheme, which includes a linear dilution of LL37 in columns 7 and 12 followed by two separate 2/3 dilution series of the cells and LL37 on columns 12 to 8 and 7 to 1. (B) Each well represents a different combination of LL37 and cell densities from which we can extract the MIC as a function of inoculum size by monitoring growth of the culture in individual wells. The solid data points refer to the wells with growing culture and the size of the marker refers to the number of repeated trial outcomes that resulted in growing cultures. The empty points refer to wells with no visible growth. A theoretical model developed later in this work nicely 1t the average MIC. (C) The growth of the cultures were monitored by an automated plate reader in terms of OD_600_. Growing cultures reach a yield comparable to each other while non-growing cultures do not exhibit consistent increase in OD_600_. Data are from column 11 of Fig 1AB and they follow the same color coding. (D) Analysis of the growth in sub-MIC cultures reveal that growth is delayed depending on the LL37 concentration, but the exponential growth rate shows no considerable change. Data follow the same color coding as Fig. 1AB. (E) Through colorimetric measurement of the concentration of a fluorescently tagged analogue of LL37 (5-FAM-LC-LL37), we can quantify the amount of peptides absorbed by *E. coli* cells. (F) The amount of absorbed 5-FAM-LC-LL37 increases with inoculum size in a culture with initial peptide concentration of 14 μM.

Each well on the microplate corresponds to a unique combination of LL37 and cell densities (Fig. 1B). The growth of the cultures in wells were monitored for 24 hours by an automated plate reader (EPOCH 2, BioTek) in terms of optical density at 600nm wavelength (OD_600_), while the plate was incubated with orbital shaking at 37°C. Growth or inhibition of growth in each well is evidently distinguishable: growing cultures reach a yield comparable to each other^1^, but the non-growing cultures do not show any consistent increase in OD_600_ over the course of the experiments except for minimal random fluctuations (Fig. 1C).

The results, averaged over four similar trials, demonstrate a distinct increase of the MIC as a function of inoculum size (Fig. 1B). (Detailed data presented in the SI Figs. S3 and S4.) The solid data points in Fig. 1B refer to the wells with a growing culture and the size of the marker refers to the number of repeated trial outcomes that resulted in growing cultures. A theoretical model developed later in this work nicely fits the average MIC.

### Sub-MIC cultures exhibit delayed growth, not slow growth

An interesting feature evidenced in the results was the extended lag phase, up to several hours, introduced by the sub-MIC concentrations of LL37 (see Fig. 1C). Despite such a ‘growth delay,’ the average doubling time of the cells (*T_D_*) did not change significantly, remaining under 30 minutes in most cases (see Fig. 1D). We tested and confirmed the stability of peptides over the duration of the experiment (Fig. S5). Hence, we hypothesized that this behavior was attributed to heterogenous cell death, where the growth of a *fraction* of the cells is inhibited, while the rest of the cells recover the normal population growth after a time delay that is correlated to the number of dead cells. This hypothesis is investigated further at the single-cell level.

### *E. coli* cells absorb and retain peptides

In the microplate experiments, direct cell-to-cell interactions are minimal as the cells are on average over 50 μm apart from each other (corresponding to inoculum size in Fig. 1AB). All the electrostatic interactions are also completely shielded^2^. We asked whether the inoculum effect is due to the absorption of peptides into the cells Clark et al. (2009). AMP absorption into bacteria has been previously discussed and quantified using various techniques. Different prokaryotes were reported to absorb 1–20 × 10^7^ AMPs/cell Steiner et al. (1988); Savini et al. (2018); Starr et al. (2016); Tran et al. (2002); Melo et al. (2011); Roversi et al. (2014), which is high enough to initiate the inoculum effect. Here we also quantified the absorption of a dye-tagged analogue of LL37 (5-FAM-LC-LL37, AnaSpec) by colorimetric measurement of the 14 μM of peptide concentration before and after incubation with *E. coli* and separation by centrifugation (see Fig. 1EF and Fig. S6 for details). We observed a reduction in peptide concentration proportional to the inoculum size with an average rate of 7.6 ± 2.1 × 10^8^AMPs/cell.

### Single-cell data demonstrate absorption and retention of peptides in target cells

We further investigated peptide absorption by tracking dye-tagged peptide action on live cells. To this end, we brought *E. coli* cells from an exponential culture to an imaging platform where they were treated with an above-MIC concentration (10 μM) of 5-FAM-LC-LL37 under agarose gel containing RDM growth media. (see Fig. S7 for the imaging platform).

We closely monitored cell growth and distribution/localization of peptides by phase contrast and 2uorescent time-lapse microscopy (Fig. 2A). By analyzing 383 cells, we observed that the inhibition of growth is followed by a rapid translocation of peptides into target cells, as quantified by a jump in the cell’s fluorescent signal (Fig. 2B). As a result, fluorescent signals showed a bimodal distribution over the course of the experiment. A large degree of temporal, cell-to-cell heterogeneity was also observed as the peptide translocation time varied for about 30 minutes for different cells (Fig. 2B). The fluorescent signal remained high after translocation confirming retention of peptides in the cells.

**Figure 2.**
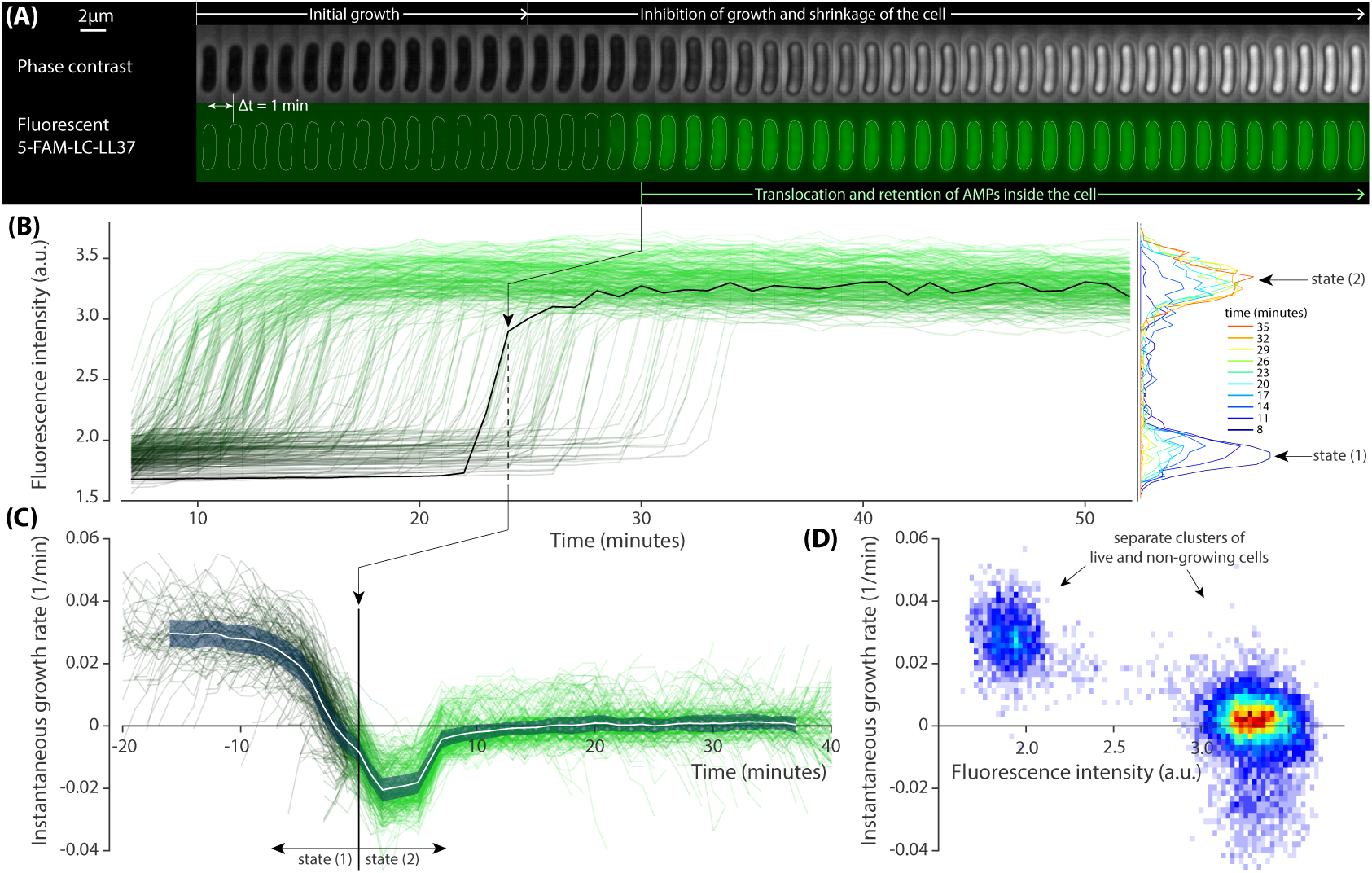
Growth inhibition of *E. coli* cells by dye-tagged LL37. (A) Sample phase contrast and fluorescent time-lapse images of an *E. coli* cell treated with a lethal dosage of the dye-tagged LL37 (5-FAM-LC-LL37). Phase contrast images show inhibition of growth and slight shrinkage of the cell after a brief growth period. The fluorescent channel shows distribution, translocation and retention of the peptides. (B) Abrupt transition in the fluorescent signal of 383 cells that occur over a span of more than 30 minutes. (C) The instantaneous growth rate of individual cells collapse on each other when plotted in referenced to the peptide translocation point. The average behavior (white line) of the collapse shows that the translocation happens shortly after the inhibition of growth and shrinkage of the cell. The shaded area denotes the standard deviation. (D) The two dimensional distribution of instantaneous growth rate and fluorescent signal depict well-separated clusters referring to a binary response of the cells to the peptides.

The instantaneous growth rate of individual cells was non-monotonic and collapsed onto each other once plotted with reference to peptide translocation time (Fig. 2C). There is a drop to negative values, indicating the shrinking of cells, which was found to be synchronized with the uptake of peptides. The growth rate reaches a steady value of zero in ~10 minutes.

As a whole, *E. coli* cells exhibited a binary physiological state over the course of the peptide action in terms of growth rate and peptide uptake. That is, the cells were found to be in either of these distinct states: (1) growing, with no significant peptide uptake; and (2) non-growing, followed by an abrupt peptide uptake. This is quantitatively evident in the scatter plot of the instantaneous growth rate as a function of fluorescence intensity, where cells segregate into two separate clusters (Fig. 2D). At the peptide concentration of 10 μM (above the MIC) all cells were initially in state (1) and then transitioned to state (2) within one generation.

### Growth inhibition is heterogeneous in sub-MIC cultures

Population-level data from microplates were suggestive, but not conclusive, of a heterogeneous growth inhibition in cultures with a sub-MIC concentration of AMPs. Hence, we proceeded with single-cell experiments as noted above with a sole modification of using a lower, sub-MIC concentration of 5-FAM-LC-LL37 (4.0 μM for the dye-tagged peptide). As such, most individual cells grew to form micro-colonies.

The striking observation was the phenotypic heterogeneity in the isogenic population of cells in each micro-colony (Fig. 3A). As a colony expanded, growth of some cells was inhibited and peptides translocated into them. we observed a similar transition from state (1) to state (2) as previously seen in above MIC cultures (Fig. 2). The difference is that at sub-MIC the transition occurred for only a fraction of the cells.

**Figure 3.**
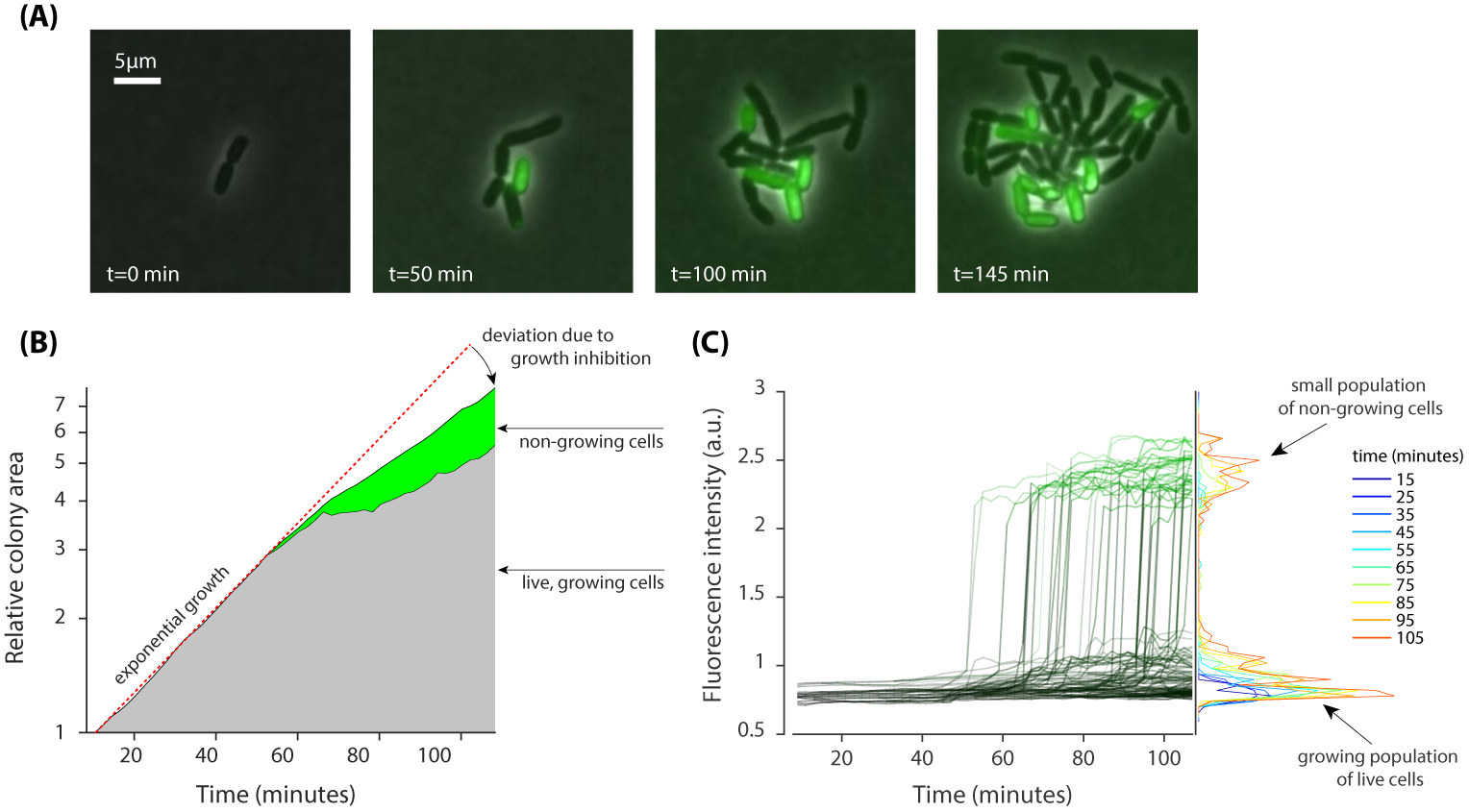
Heterogenous growth inhibition and peptide absorption at a sub-MIC concentration of peptides. (A) Sample time-lapse images (overlay of phase contrast and fluorescent image) of dividing *E. coli* cells show that growth of only some cells is inhibited in a growing colony. Phase contrast images show growth of a micro-colony and the fluorescent channel (green) shows translocation of the dye-tagged LL37 (5-FAM-LC-LL37) in the cells whose growth is inhibited. (B) Relative growth of the total area of 13 separate colonies, consisting of 280 cells, depicts initial exponential expansion (grey area) until the appearance of non-growing subpopulation (green area). (C) An abrupt transition in fluorescent signal is observed when growth is inhibited in cells. The transition for individual cells is similar to that in above MIC cultures (see Fig. 2).

Analysis of 13 separate micro-colonies, consisting of a total of 280 cells (over the course of the experiment), showed that the relative size of the colonies initially increased exponentially until the appearance of non-growing cells (Fig. 3B). The fluorescence intensity of non-growing cells showed an abrupt transition, as in the case of above MIC cultures, with comparable relative changes in the signal, which suggests that a similar number of peptides are taken by each cell (Fig. 3C).

### Mathematical model based on peptide absorption reproduces and explains exper-imental observations

In order to test whether the absorption of peptides can explain the inoculum effect, we developed a mathematical model with minimal single-cell assumptions (Fig. 4A), which describes the time evolution of the mean concentration [B] of growing bacteria and the mean concentration [P] of free AMPs in the solution. The model describes two processes: (1) bacteria are assumed to divide with a constant rate *k_D_* = ln 2/*T_D_*, where *T_D_* ≈ 23min is the average doubling time; (2) AMPs kill growing bacteria with a rate *k_k_*, and afterwards each dead cell quickly takes up *N* AMPs (see Figs. 2B and 3C). These AMPs are bound to the membrane as well as to the cytoplasm of the cell and are not recycle to attack other cells. The time evolution of concentrations of bacteria [B] and available AMPs [P] is described by the following equations:

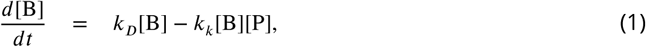

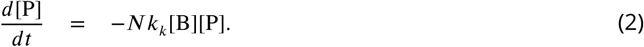

**Figure 4.**
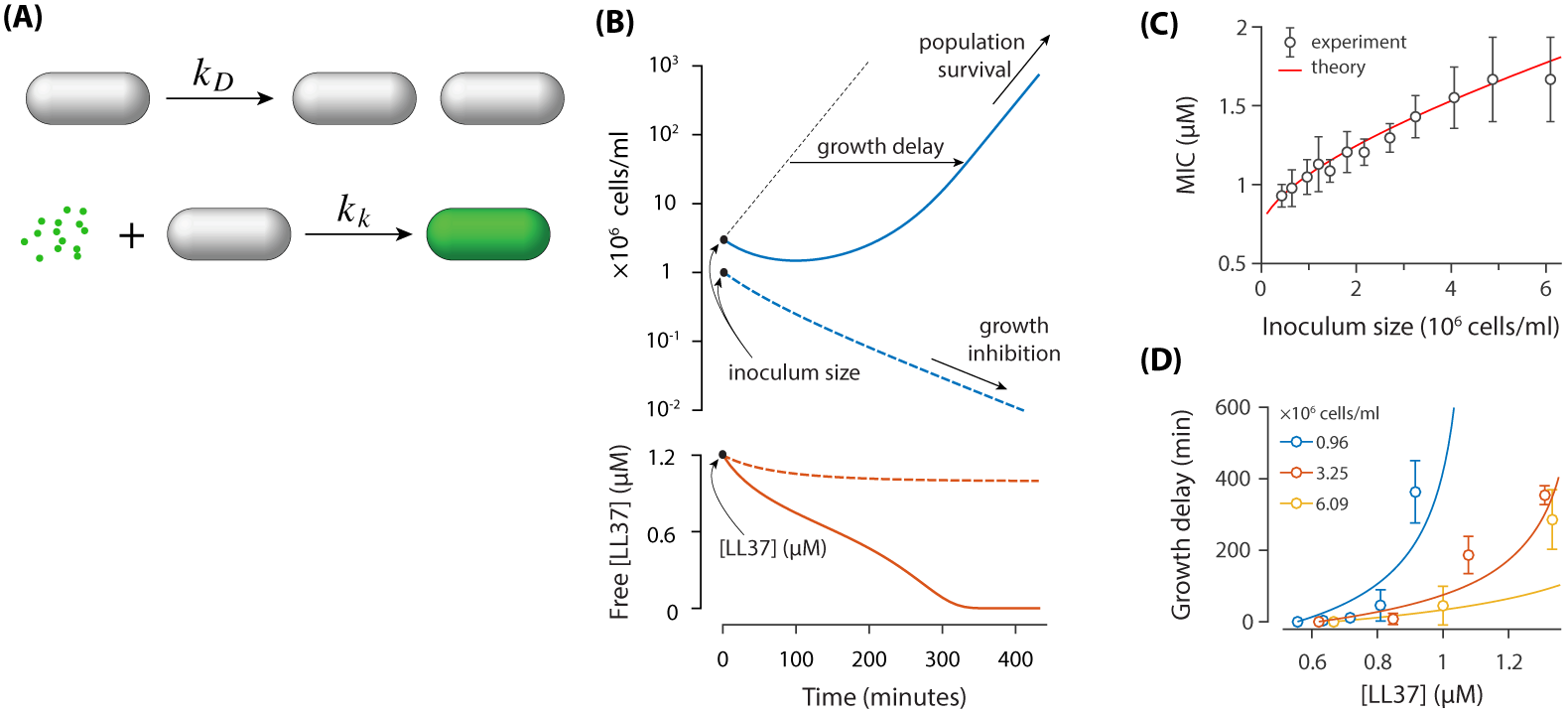
A theoretical model based on the absorption of peptides in *E. coli* cells. (A) *E. coli* cells replicate with a rate of *k_D_* and they get killed with a rate of *k_k_*. Each dead cell quickly absorbs *N* AMPs. (B) Demonstration of the inoculum effect for the initial AMP concentration [LL37]=1.2 μM. A culture with high inoculum size (3 × 10^6^ cells/ml) survives as peptides deplete (solid lines) whereas growth in small inoculum size (10^6^ cells/ml) is inhibited with excess peptides remaining in the solution (dashed lines). Despite the survival of culture with high inoculum size, the growth is delayed. (C) Comparison of the MIC between the theoretical model and the experimental data. (D) Growth delay as a function of [LL37]. Solid lines are theoretical results (not a 1t) and circles represent experimental data from microplate experiments (Fig. 1). The delay is calculated with respect to the lowest AMP concentration (row H) of Fig. 1AB.

This model predicts two different outcomes (see Fig. 4B) depending on the initial concentrations of bacteria and AMPs: (1) The population of bacteria goes extinct for a sufficiently large concentration of AMPs, i.e. above MIC; (2) The population of bacteria can recover in a low concentration of AMPs, i.e. below MIC. The two unknown parameters of the model were fitted to best approximate the MIC dependence on the inoculum size (Fig. 4C). The fit resulted in a killing rate *k_k_* = 0.040*μ*M^−1^ min^−1^ and *N* = 3.8 × 10^7^ AMPs absorbed per dead cell. Note that in the limit, where the initial concentration of bacteria goes to zero, the MIC value approaches the finite value *k_D_*/*k_k_* = 0.75*μ*M [see Eq. (1)]. This is consistent with a previous model by Stella group Savini et al. (2017), which considered that bacteria get killed once the number of peptides bound to cell membrane reaches a certain threshold (Note that the number of surface bound peptides correlates with the concentration of free peptides in solution).

We further examined whether the model can reproduce other experimental data without additional fitting. In particular we tested whether the model could predict the growth delay in surviving bacterial population when the concentration of AMPs is increased (see Fig. 1C). The predictions of our model agree reasonably well with experimental results for the growth delay of population (see Fig. 4D), given the simplicity of the model.

### The action of dye-tagged LL37 is cell-cycle dependent

The temporal heterogeneity we reported in growth inhibition and peptide retention is a key factor for the emergence of the surviving subpopulation. The wide distribution of ~30 minutes (above MIC cultures, Fig. 2B) is puzzling, as all the cells experienced the same environmental conditions. We looked at the correlations of the translocation time with two cell size measures to investigate any dependence on the physiological conditions of the target cells. A strong negative correlation was observed between peptide translocation time and initial cell length, indicating that small cells can resist AMPs’ action, growing until a later time point (Fig. 5 left panel). In contrast, cell length at translocation time did not show any correlation with the translocation time (Fig. 5 right panel). This clearly demonstrated that the action of 5-FAM-LC-LL37 is cell-cycle and cell-age dependent. This seems in agreement with the findings made by the Weisshaar lab, where LL37 peptides were observed to first bind to the septum of dividing cells. Thus, the higher chance to act on larger, dividing cells, as opposed to small growing cells Sochacki et al. (2011). The stronger binding of LL37 to the septum area is not well understood but may have multiple physical origins. Among various possibilities, the Wong lab has shown LL37 to preferentially bind to membranes with negative Gaus-sian curvature, a geometry that can be found in the septum of rod shaped microorganisms Yang et al. (2008); Schmidt et al. (2011).

**Figure 5.**
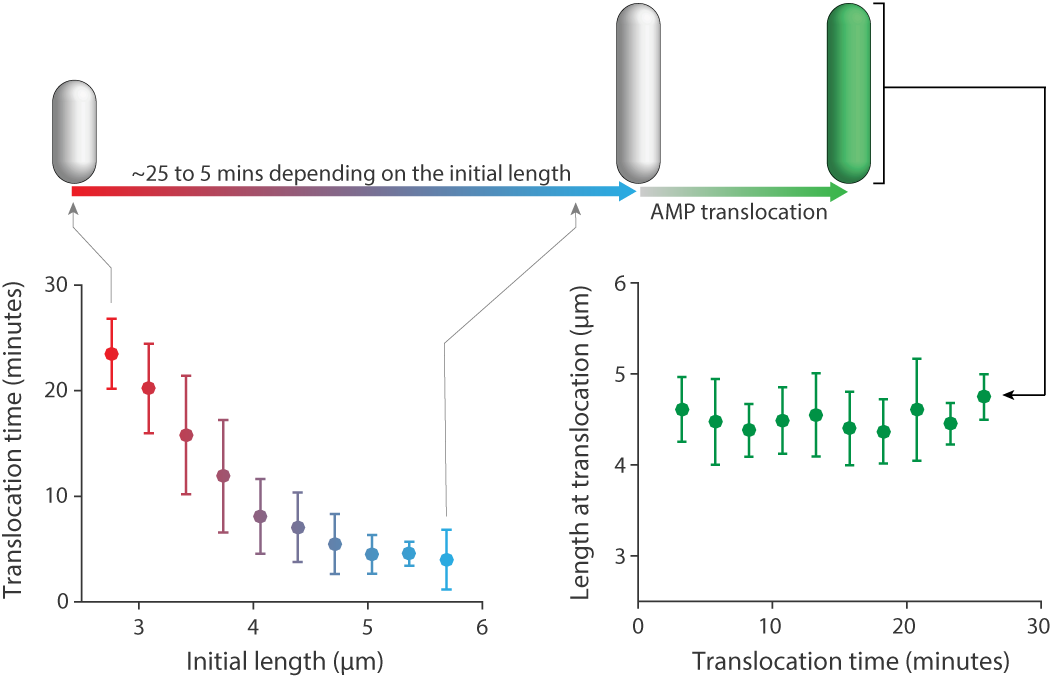
Correlations between the peptide translocation time and cell size. (Left panel) A strong negative correlation between peptide translocation time and initial cell length indicates resistance of small cells to peptides until a later time point. (Right panel) Cell length at translocation time and the time of translocation is not correlated.

## Discussion

Heterogeneities in bacterial response to antibiotics can be critical if leading to the survival of a subpopulation that can recover population growth Coates et al. (2018). In this work, we discovered an unexpected absorption and retention of an antimicrobial peptide (LL37 and the dye-tagged analogue, 5-FAM-LC-LL37) in *E. coli* cells, which under sub-MIC concentrations led to the emergence of two distinct subpopulations in an isogenic bacterial culture: a group of cells that retain peptides after their growth is inhibited and a group of surviving cells that grow owing to the reduction of the “free” peptide concentration by the other group. This “passive cooperation” is an interesting feature of an *E. coli* culture where cells do not have any form of active communication, unlike ion-channel based cooperation in *Bacillus subtilis* biofilms Prindle et al. (2015); Liu et al. (2017, 2015).

At the cellular and molecular scales, a distinct feature of the AMP’s mechanism of action is its collective nature, where a large number of AMPs are required to first bind to and then disrupt the cell membranes to kill the cell. AMP’s absorption has been discussed and quantified previously for different microorganisms and peptides using various techniques Steiner et al. (1988); Savini et al. (2018); Starr et al. (2016); Tran et al. (2002); Melo et al. (2011); Roversi et al. (2014). Utilizing live, single-cell microscopy, we observed the temporal and cell-to-cell heterogeneity in the peptide absorption into *E. coli* cells, which goes beyond the membrane binding.

The integration of the population and single-cell data, combined with the theoretical modeling, presented in this work, provides strong evidence that the inoculum effect for the LL37 arises from the retention of LL37 in target cells. Despite the seeming complexity of the LL37 partitioning in a population of bacterial cells, the picture at the single-cell level is simple and binary, consistent in cultures with above MIC and sub MIC concentrations of AMPs: occurrence of growth inhibition depends on the free peptide concentration and is followed by an abrupt, permanent translocation of peptides into the target cells.

Our quantification of peptide retention (~3.8 × 10^7^ peptides/cell) is within the range reported using various methods Steiner et al. (1988); Savini et al. (2018); Starr et al. (2016); Tran et al. (2002); Melo et al. (2011); Roversi et al. (2014). Yet, this large number raises the question of which molecules inside of the cells are interacting with LL37. The negative charge of DNA as well as some proteins can provide binding sites for LL37. To categorically distinguish between these two, we utilized an *E. coli* strain lacking the septum positioning *minCDE* system, which produces enucleated mini-cells (Fig. 6A). The mini-cells do not contain DNA as confirmed by the localization of a fluorescently tagged histone-like protein *hupA* (Fig. 6A, see SI for the details of the strains genotype)^3^. Translocation of 5-FAM-LC-LL37 showed absorption and retention into mini-cells in a similar, qualitative fashion as seen in regular cells (Fig. 6B), suggesting the presence of significant peptide-protein interactions.

**Figure 6.**
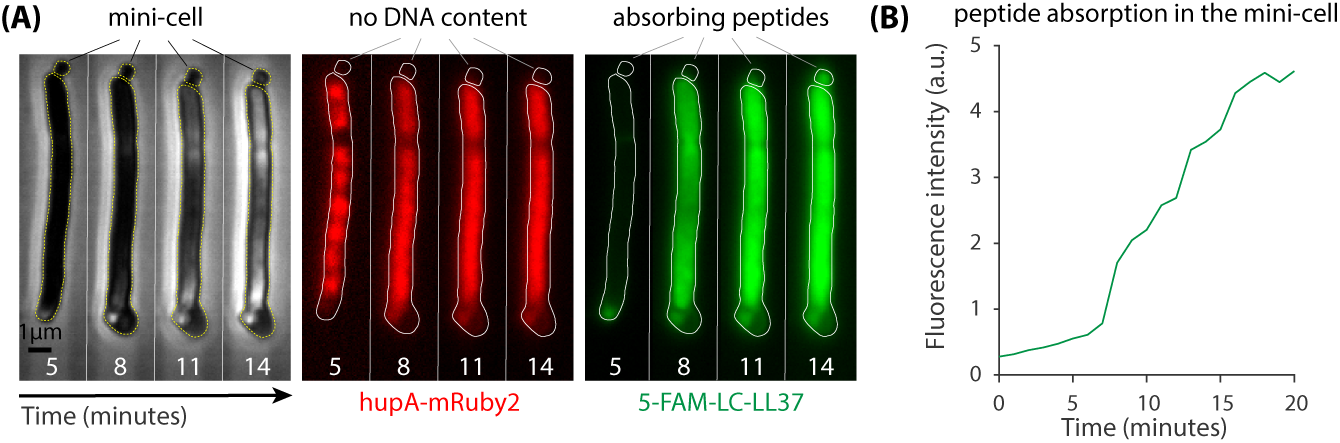
∆*minCDE* strain of *E. coli* used to test significance of peptide-protein interactions. (A) Lack of septum positioning system produces enucleated mini-cells (left panel) that do not contain DNA as indicated by the localization of *hupA-mRuby2* protein (middle panel). Mini-cells absorb a large amount of 5-FAM-LC-LL37 peptides similar to wild-type cells (right panel) (B) Average fluorescence intensity of a mini-cell as a function of time suggests a significant peptide interactions with proteins in the mini-cells.

While our results provide a quantitative picture of LL37 partitioning and acting in an *E. coli* population, they open new questions on the molecular and evolutionary basis of their activity. First, the strong absorption of peptides into the cytoplasmic area raises questions on the nature and impact of this intercellular binding: what specific proteins and domains are peptides binding to? How does the peptide binding perturb the protein functions? Second, the population survivability as a result of peptide absorption raises questions on the evolutionary dynamics of this phenomena: how do *E. coli* cells evolve to achieve this cooperative fit? How does this phenomena affect multi-species cultures with prokaryotic and eukaryotic organisms?

Finally, our findings imply an important dynamic for the activity of LL37 (and possibly other AMPs) in the host, multicellular systems. Considering the peptide absorption into the target cells, AMP concentration should not be assumed the only key factor, but the rate of the expression of the AMPs by the host is also decisive in determining effectivity of AMPs. The expression rate competes with the rate of absorption of the AMPs in the bacterial cells. In the results presented in this work, we focused on ‘closed’ systems where the total number of AMPs remained constant. As a future direction, one could examine growth inhibition of bacterial cultures experiencing an influx of the AMPs.

## Methods and Materials

### Bacterial Strains and Growth Conditions

In this work, we used derivatives of a prototrophic *Escherichia coli* K12 strain, NCM3722, that was constructed, sequenced, and extensively tested by Kustu and Jun labs Soupene et al. (2003); Brown and Jun (2015). In all microplate and single-cell experiments, we used ST08, a nonmotile derivative of NCM3722 (∆motA) (a gift from Suckjoon Jun’s Lab at the University of California, San Diego), except for experiments with mini-cell producing strains where we used ST20. This strain possesses a deficiency in septum positioning system (∆minCDE) and a DNA marker (*hupA*-*mRuby*2). ST20 was constructed using standard P1 transduction to transfer the gene deletion minCDE::aph from PAL40 (from Petra Levin’s Lab at the Washington University, St. Louise) to ST12, a construct from ST08 with the infusion of *mRuby*2 fluorescent protein with the histone like protein *hupA* (from Suckjoon Jun’s Lab).

In all experiments, a MOPS based rich defined media (RDM) was used, developed by Fred Neidhardt Neidhardt et al. (1974), which is commercially available from Teknova Inc. The average generation time of the *E. coli* strain used in this study was 23 minutes in RDM at 37°C. All cells in the experimental samples were grown to early or mid log phase prior to the start of the experiment in a 37°C water bath shaker, set to 240 rpm.

### Live-cell Imaging Platform

To monitor the inhibition of growth for *E. coli* cells by antimicrobial peptides in a live microscopy setting we chose to use agarose pads to immobilize them. Inspired by previous works Moffitt et al. (2012); Priest et al. (2017), we developed a system for patterning and housing specific amounts of agarose gel suitable for long term microscopy needs. The patterns are parallel channels that allow cells to spread and move away from one another under the agarose pad, while aligned in certain directions to help with the image analysis and cell segmentation. The housing also reduces the chance of evaporation of the liquid culture, thus allowing the gel to be used for hours at 37°C during microscopy.

### Phase Contrast and Fluorescent Microscopy

An inverted microscope (Nikon Ti-E) equipped with the Perfect Focus System (PFS 3), a 100x oil immersion objective lens (NA 1.45), and an Andor Zyla sCMOS camera were used for imaging. The light source that was used for the phase contrast microscopy was made possible with the help of LED transmission light (TLED, Sutter Instruments 400-700nm) and Spectra X light engine (Lumencor), which was used for fluorescent imaging.

The illumination condition for phase contrast was 50ms exposure with an illumination intensity set to 10% of the max TLED intensity. The fluorescent images for the 5-FAM dye were taken with the excitation wavelength of 485 while using a quad band filter (DAPI/FITC/TRITC/Cy5, 84000, Chroma Technologies).

### Analysis of the Theoretical Model with Mathematica

Differential equations [Eqs. (1-2)] describing the population dynamics model were analyzed in Mathematica with the function *NDSolve*. For each initial concentration of bacteria and peptides, the evolution of both populations were analyzed during the subsequent *T*_max_ = 1000 min. If the concentration of peptides [P(*T*_max_)] at the end of simulation was smaller than *k_D_*/*k_k_*, then the bacterial population survived [see Eqs. (1-2)]. On the other hand, if the concentration of peptides [P(*T* max)] *> k_D_*/*k_k_*, then the bacterial population most likely went extinct. The MIC concentration for a given initial concentration of bacteria was obtained by requiring [P(*T* max)] = *k_D_*/*k_k_*, which was found using the bisection method. Finally, the unknown model parameters *k_k_* and *N* were obtained by minimizing the error between the values of MIC from the model and the experimental data (see Fig. 4C) by using the function *FindMinimum*.

## Acknowledgments

We acknowledge funding support from the National Institute of Health grant 1R15GM124640, the Pilot Project grant under 1RL5GM118975, and the National Science Foundation (NSF) grants Research Experiences for Undergraduates (REU) site grant (EEC-1559973), Partnership in Research and Education in Materials (PREM) between the W. M. Keck Computational Materials Theory Center (CMTC) at California State University, Northridge, and Princeton Center for Complex Materials (PCCM) (DMR-1205734), and the Materials Research Science and Engineering Center Program through the PCCM (DMR-1420541).

## Footnotes

1 Yield in microplate experiments are limited by the ‘ghost’ effect as there is no chance of dilution prior to OD measurement in dense cultures. Nevertheless, for the results in this work, yield is not used to quantify growth of the culture.

2 The Debye screening length in RDM is 12Å with 61mM of monovalent and 0.5mM of divalent ions.

3 The mini-cells are not viable due to a lack of DNA, whereas the other sister cells are longer than average, containing more than one round segregated DNA. Despite the abnormal cell size, the DNA-containing sisters are mostly viable.

